# Neuronal membrane proteasomes homeostatically regulate neural circuit activity *in vivo* and are required for learning-induced behavioral plasticity

**DOI:** 10.1101/2022.01.29.478314

**Authors:** Hai-yan He, Kapil Ramachandran, Natalie McLain, Regina Faulkner, Arifa Ahsan, Reshmi Bera, Seth S Margolis, Hollis T Cline

## Abstract

Protein degradation is critical for brain function through processes that remain poorly understood. Here we investigated the *in vivo* function of a recently reported neuronal membrane-associated proteasome (NMP) in the brain of *Xenopus laevis* tadpoles. We demonstrated that NMPs are present in the tadpole brain with biochemistry and electron microscopy, and showed that they actively degrade neuronal activity-induced nascent proteins. Using *in vivo* calcium imaging in the optic tectum, we showed that acute inhibition of NMP function rapidly increased spontaneous neuronal activity, resulting in hyper-synchronization among tectal neurons. At the circuit level, inhibiting NMPs abolished learning-dependent improvement in a visuomotor behavior paradigm in live animals. Our data provide the first *in vivo* characterization of NMP functions in the vertebrate nervous system and suggest that NMP-mediated degradation of activity-induced nascent proteins may serve as a homeostatic modulatory mechanism in neurons that is critical for regulating neuronal activity and experience-dependent circuit plasticity.

## INTRODUCTION

Proteostasis, the collective protein synthesis, folding and degradation mechanisms that maintain the integrity of the proteome, is pivotal for the health and normal function of the nervous system^1^. Both protein synthesis and protein degradation have been shown to be necessary for synaptic plasticity and homeostasis^2–13^. As one of the two major proteolysis pathways that have been discovered to be employed by neurons, proteasomes have been widely studied.

Proteasomes are large macromolecular complexes with 28 subunits and six catalytic subunits^14–17^. Together, these make up the core 20S particle (CP) which can interact with a set of proteins that make up a 19S cap, forming the 26S (singly-capped) or 30S (doubly-capped) proteasome. 19S capped proteasomes are responsible for the majority of ubiquitin-dependent degradation in the nervous system and require ATP. 20S uncapped proteasomes do not require ubiquitin or ATP and have been shown to operate independently to degrade intracellular proteins, such as damaged/oxidized or unstructured proteins^18–20^. Emerging experimental evidence suggests a significant proportion of 20S uncapped proteasome exists in the brain and may play critical roles in neuronal physiology^1, 21–23^.

Recent work identified an uncapped 20S proteasome complex that is tightly associated with the neuronal plasma membrane in the mammalian central nervous system (CNS)^22, 23^. Blocking the catalytic activity of this neuronal membrane proteasome (NMP) in cortical neuronal cultures rapidly altered neuronal activity under pharmacological stimulation^23^. Furthermore, NMPs function at least in part through ubiquitin independent degradation of activity-induced nascent polypeptides as they are actively translated^22^, suggesting that NMPs could play as yet undiscovered roles in regulating neuronal activity and activity-dependent proteostasis. The potential functions of NMPs in living animals has yet to be described.

Activity-induced protein synthesis is a signature of associative synaptic plasticity^24–27^. These newly synthesized proteins are thought to play important roles in the expression and maintenance of the activity-induced plasticity. For example, blocking protein synthesis during LTP induction obliterates late-phase LTP^28, 29^. On the other hand, blocking protein degradation by blocking proteasomes also prevents the long-term expression of synaptic plasticity^7, 11, 30–32^. Although it is generally recognized that the ubiquitin-dependent proteasome pathway plays a major role in these processes^6, 11, 33^, many experiments used inhibitors targeted to the 20S core particle of proteasomes^34^, which would also inhibit NMPs^23^. Taken together, we hypothesized that NMPs are involved in the regulation of the proteostasis of activity-induced nascent proteins that is important for neuronal and circuit function and behavioral plasticity in intact neural circuits. We tested our hypothesis in the *Xenopus laevis* tadpole visual system. The visual circuit of tadpoles is a robust experimental system for investigation of experience-dependent plasticity^35^, and studies of the tadpole visual circuit have contributed significantly to our understanding of different mechanisms underlying plasticity (for reviews see ^35–37^). Prior studies in Xenopus visual system have applied bio-orthogonal non-canonical amino acid tagging (BONCAT) labeling of nascent proteins in living tadpole brain to identify visual experience-induced nascent proteins and demonstrate the requirement of nascent proteins for visual experience-dependent behavioral plasticity ^38–41^. This system is thus well suited to test NMP function through *in vivo* quantitative analysis of proteostasis of BONCAT-labeled nascent proteins, analysis of neuronal activity in the intact visual circuit and behavioral evaluation of circuit function and plasticity.

Here we examined the proteolytic function of NMPs in the brains of live tadpoles. We found that the core proteasome subunits (α1-7) are abundantly expressed in the tadpole brain, and are associated with the neuronal membrane, as shown by biochemistry and EM studies. Using BONCAT labeling of nascent proteins in live tadpole brain^38–41^, we showed that inhibiting NMPs significantly increased nascent protein levels, especially in response to increased neuronal activity. This coincided with a rapid increase in spontaneous neuronal activity and an increase in synchronous activity across the neural circuit of intact animals. Considering the extensive evidence that aberrant synchronization in brain circuits disrupts cognitive function and behavior^42–, 46^, these data suggest that the role NMPs play in regulating neuronal activity in the intact brain may be particularly important for maintaining circuit activity within a physiologically relevant range. Indeed, we found that inhibiting NMP activity abolished visual experience-dependent behavioral plasticity. Our results demonstrate that NMPs are functionally active *in vivo* and identify a role for NMPs in modulating activity-dependent neuronal proteostasis and experience-dependent circuit and behavioral plasticity.

## RESULTS

### NMPs are present in neurons in tadpole brains

To study NMP functions *in vivo*, we employed a membrane impermeable 20S core proteasome inhibitor, Biotin-Epoxomicin (BE). BE is the biotinylated form of Epoxomicin (Epox), a highly specific proteasome inhibitor which covalently binds to catalytic b subunits of the core 20S proteasome^47^. Biotinylation renders BE impermeability to cell membrane. When BE is added to live neuronal cultures, it specifically binds to and blocks NMPs on the neuronal membrane^22, 23^. Therefore, BE can be used both as a marker and as a highly specific inhibitor for NMPs in living neurons. To determine if 20S core subunits of proteasomes are present in the neuronal membrane, we injected BE (1 mM) into brain ventricles of live tadpoles and used immunohistochemistry to examine the distribution of BE-bound NMPs in the optic tectum (Figure 1A) with anti-biotin antibody. Strong biotin signal was observed in the neuronal soma layer and the neuropil layer of the optic tectum from BE-injected animals, with little signal seen in the ventricular layer where the neuroprogenitor cells reside (Figure 1B). No signal was seen in un- injected brain sections that were processed and imaged under the same conditions. These data suggested that BE-bound 20S core particles (CP) are not only present in the tadpole brain but also are preferentially associated with neurons, supporting the presence of NMPs in tectal neurons. To further confirm that BE specifically binds to the proteasome core subunits in the neuronal membrane in tadpole brain, we harvested tadpole brain tissue from BE or epoxomicin-injected animals as well as un-injected controls, and performed western blots for NMP subunits in cytosolic and membrane preparations. Epoxomicin-injected and un-injected animals served as a negative control for biotin. In addition, to assess the timeframe over which BE remains bound to NMPs *in vivo*, we collected brain tissue either 30 min or 6 hr after the injection. As seen in both injected and un-injected samples, the 20S proteasome core subunits (α1-7) are abundantly expressed in the tadpole brain and are present in the isolated membrane fraction (Fig 1C). Importantly, the biotin signal was only observed in the membrane fraction in BE-injected samples and appeared at the molecular weight corresponding to the *β* subunits (20-25 kD). In addition, the biotin signal from BE is detected in bands corresponding to the molecular weight of the 20S *β* subunits in the membrane preparation both at 30 minutes and 6hr after the injection but not in the cytosolic fraction, suggesting that BE is irreversibly bound to the core subunits on the membrane, consistent with prior work demonstrating epoxomicin binding to the core subunits^22, 23, 47^. To further confirm the binding specificity of BE, we preinjected tadpoles with epoxomicin before intraventricular injection of BE, and used neutravidin-coated resin to pull down biotin-bound proteins from total brain lysate. Western blotting of the neutravidin-pulldown samples showed that 20S *β*5 subunits were co-precipitated with biotin in the BE-injected samples, but not in either Epoxomicin-preinjected or un-injected samples (Figure 1D). Labeling of the BE-specific biotin bands corresponding to the 20S *β* subunits was also blocked in the epoxomicin-preinjected samples. The occlusion of BE-binding to the 20S *β* subunits confirmed that BE and epoxomicin share binding sites. We also labeled the western blots with antibodies to Actin, a cytosolic protein, and GluA1, which is known to enrich in the neuronal membrane, to serve as quality controls for the membrane fractionation preparation. Taken together, these data demonstrate the existence of NMPs in *Xenopus laevis* brain, indicating that NMPs are conserved in non-mammalian vertebrates.

**Figure 1.**
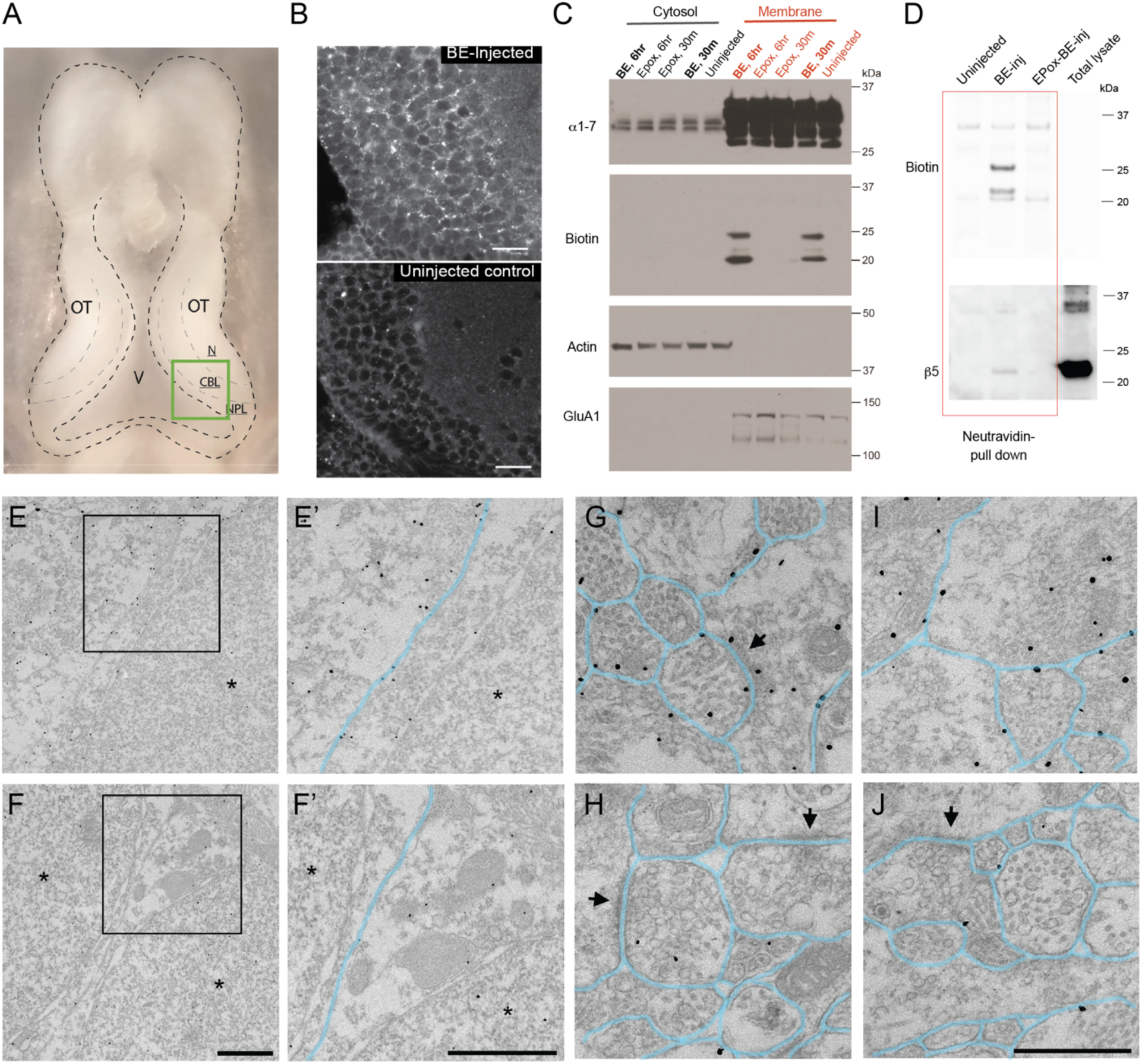
NMPs are present in tadpole neurons. A. Anatomical illustration of cellular organization of the optic tectum (OT) and the ventricle (V). CBL: neuronal cell body layer. NPL: neural progenitor cell layer. N: Neuropil. B. Immunohistological localization of NMPs bound to biotin-Epoxomicin (BE). Vibratome sections of optic tectum immuno-stained with anti-biotin antibody show punctate biotin signal in neuronal cell body layers and the neuropil in BE injected tectum but not un-injected control. Scale bar = 20μm. C. Biochemical evidence for NMPs in tadpole brain. Tadpole brain tissue was collected at 30m or 6hr after either BE or epoxomicin (EPox) injection and equivalent protein amounts from membrane and cytosolic fractions were labeled with indicated antibodies on western blots. The core proteasome subunits (α1-7) are abundantly expressed in the tadpole brain and are enriched in the membrane fraction. Importantly, the biotin signal was recovered at the molecular weight corresponding to the 20S catalytic subunits bound to the injected BE in (and only in) the membrane preparation both at 30 minutes and 6hr after the injection. Actin, a cytosolic protein, and GluA1, which is enriched in the neuronal membrane, serve as positive controls for the cell fractionation prep. D. Pre-injection of epoxomicin occluded BE-binding to NMPs. Brain lysates from un-injected, BE-injected, and EPox pre-injected (Epox-BE-inj) samples were purified by neutravidin pull down to co-precipitate BE-bound proteins and blotted with biotin and 20S *β*5 antibody. E-J. Ultrastructural distribution of the 20S proteasome shown by pre-embedding immunogold labeling with anti-*α*1-7 20S proteasome subunit antibody and FluoroNanogold secondary antibody (1.4nm-diameter gold particles). Plasma membranes are highlighted in light blue. (E, F) Ultramicrographs from the tectal cell body layer showing anti- *α*1-7 20S immunogold labeling (E) and no primary controls (F). Boxed regions are shown at higher magnification in (E’, F’). Immunogold particles identifying 20S *α* subunits are associated with the plasma and nuclear membranes and found in the cytoplasm. Asterisks mark nuclei. Scale bars = 1um. (G-J) Ultramicrographs from the tectal neuropil labeled with anti-*α*1-7 20S antibodies (G, I) or no primary control (H, J). Numerous immunogold particles identifying 20S *α* subunits are localized at or near the plasma membrane in G, I. 20S *α* subunits are also visible at synaptic sites, which are marked by arrows. Scale bar = 500nm.

To further probe the subcellular and ultrastructural localization of NMPs in the optic tectum, we employed pre-embedding immuno-EM using antibodies against the alpha subunits of the 20S CP and FluoroNanogold secondary antibody (Fig1 E, E’, FG, I). Samples without primary antibody were used as control and have relatively low background levels of Nanogold labeling (Fig1 F, F’,H, J). Ultramicrographs from both the tectal cell body layer (Fig1 E,E’) and the neuropil layer (Fig1 G, I) demonstrated that 20S alpha subunits are associated with the plasma membrane, in addition to the nuclear membrane and the cytoplasm, which are known to contain both 26S and free 20S proteasomes^20^. We also observed 20S CP immunoreactivity at synaptic sites (Fig 1G, arrows), similar to what has been reported in mouse hippocampal neurons^23^. These data support the presence of NMPs in the tadpole brain, and their localization indicates a possible role in neuronal activity dependent processes.

### NMPs actively degrade nascent proteins in the tadpole brain

Prior studies in mouse cortical neuronal cultures indicated that NMPs, independent of ubiquitin, degrade nascent proteins synthesized in response to increased neuronal activity^22^. To determine whether nascent proteins are subjected to NMP degradation *in vivo*, we measured nascent protein production from pharmacologically stimulated tectal neurons in tadpole brains, in the presence and absence of BE. For quantitative assessment of nascent protein production within a short period, we modified a previously published protocol using BONCAT^48^, that we had applied to live tadpole brains^40, 41^. Briefly, animals were intraventricularly injected with the non-canonical amino acid AHA to label nascent proteins over a period of 30min under either basal or stimulated conditions. To effectively stimulate a large population of tectal neurons, we pre-treated animals with intraventricular injection of Bicuculline Methiodide (BMI), which blocks GABAergic inhibition and significantly increases neuronal activity (Fig 2A). 50 µM BE was injected together with AHA under either basal or stimulated condition to inhibit NMPs during the 30 min labeling period. Whole brain tissue was dissected and processed with click chemistry to specifically tag AHA-labeled nascent proteins with biotin. Dot blots immunolabeled with biotin antibody were used to assess the level of nascent proteins in the protein sample. Neither endogenous biotin nor the biotin signal from the injected BE (50 µM) was detectable by dot blot under the linear exposure range for the biotinylated AHA-tagged nascent proteins, making it possibly to quantitatively monitor tagged nascent proteins (supplemental Fig 1). Under the basal condition, AHA-tagged nascent proteins were readily detected in tadpole brain tissue after the 30 minutes AHA-incubation period, consistent with previous reports in neuronal cultures^49^. In addition, treatment with the protein synthesis inhibitor, anisomycin, blocked all of the biotin signals (Fig 2B), confirming that AHA-tagged nascent proteins were the sole source for the biotin signal detected in dot blots. Consistent with prior reports of rapidly increased protein synthesis in response to heightened neuronal activity^40, 41^, the level of AHA-tagged nascent proteins increased significantly in response to BMI treatment compared to basal condition. Furthermore, inhibiting NMPs with BE in the presence of BMI induced a further significant increase in the level of nascent proteins (BMI: 1.32 ± 0.08; BMI-BE: 1.47 ± 0.07; all normalized to control from the same batch), suggesting that about a third of nascent proteins synthesized in response to increased neuronal activity was actively degraded by NMPs *in vivo*. In contrast, under basal conditions, the level of AHA-tagged nascent proteins was not significantly changed in the presence of BE (BE: 0.93 ± 0.05, normalized to control from the same batch, Fig 2B). This lack of difference could either be due to lower NMP activity level or relatively low level of protein synthesis under basal conditions.

**Figure 2.**
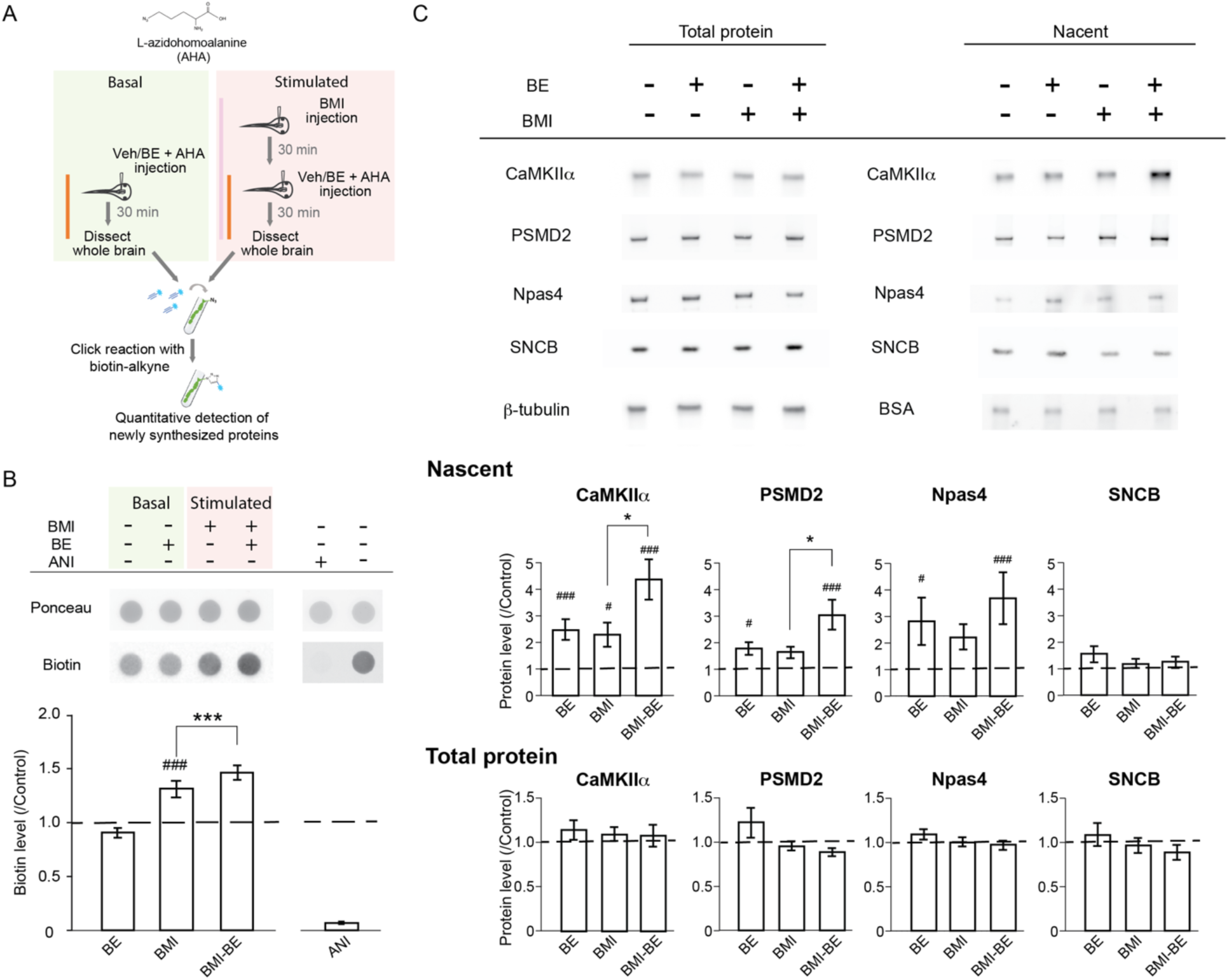
NMPs degrade nascent proteins in the tadpole brain under both basal and stimulated conditions. A. Experimental design and time line. Animals under basal or stimulated (treated with BMI) conditions were injected in the brain ventricle with either vehicle (Veh) or BE together with AHA to specifically label nascent proteins using BONCAT. B. Top: Representative dot blot with Ponceau staining and AHA-biotin immunolabeling under basal and BMI-stimulated conditions. Bottom: summary data of AHA-biotin labelled total nascent protein level detected by biotin signal in dot blots. Data from different experimental groups was normalized to the corresponding control group (marked by the dashed line) from the same batch of animals that was run on the same blot. Data shown as mean ± S.E.M, n = 24 batches. RM one-way ANOVA with Bonferroni’s multiple comparison test. ^###^: p< 0.001, comparing to control group. ***: p< 0.001, comparison as marked. Anisomycin (ANI) was co-injected with AHA under control condition in a subset (n = 3 batches) of experiments to confirm that the detected biotin-signal was from nascent proteins. C. CaMKII*α*, PSMD2, Npas4, but not SNCB are actively degraded by NMPs *in vivo*. Representative western blots (top) and quantification (bottom) for each protein, as well as loading controls (*β*-tubulin for total protein and biotinylated BSA for purified nascent proteins). Experimental conditions for each group are shown on the top. All data were normalized to the corresponding control group (dashed line) from the same batch that was run on the same blot. Friedman test with Dunn’s multiple comparison test. ^#^: p<0.05, ^###^: p< 0.001, comparing to control group. *: p<0.05, comparison as marked. n = 12 batches for Npas4, n = 16 batches for PSMD2, CaMKII*α*, SNCB.

NMP-mediated protein degradation is reportedly ubiquitin-independent^22^. To test if this is also the case *in vivo*, we used ubiquitin dot blots to assess ubiquitin levels in total protein samples from tadpole brains. As expected, no change was seen in the ubiquitin level in BE-treated samples compared to the control group under either basal or stimulated conditions (supplemental Figure 2). As a positive control, epoxomicin treatment, which inhibits all proteasome activity including the ubiquitin-dependent protein degradation, significantly increased the ubiquitin level in the brain (supplemental Figure 2).

Proteomic studies in neuronal cultures indicate that NMP substrates include some activity-induced immediate early genes (IEGs, e.g. c-fos, NPas4, and Egr1), but not others^22^, suggesting NMP-mediating degradation of nascent proteins may be substrate-selective. To examine if NMPs also exhibit substrate selectivity *in vivo*, or if they invariantly degrade all nascent proteins, we purified the biotinylated AHA-labelled nascent proteins with neutravidin-coated resin from the brain of control, BMI, and/or BE injected animals. We first verified the specificity of the neutravidin-purification of biotinylated nascent proteins with negative controls of AHA-labeled samples without click chemistry for biotinylation (supplemental Figure 3). Samples were then prepared for immunoblot analysis using antibodies raised against individual protein substrates (CaMKII, PSMD2, Npas4 and SNCB), which were selected based on prior reports of their involvement in activity-dependent protein synthesis, or the lack of it ^22, 40, 41^. CaMKII*α* was readily detected in the nascent protein samples under basal control conditions. Increasing neuronal activity with BMI treatment significantly increased nascent CaMKII*α*, consistent with prior reports of activity-induced upregulation of CaMKII*α* synthesis. Inhibiting NMPs with BE in BMI-treated brains resulted in a further significant increase in nascent CaMKII*α* (Fig 2C), suggesting that NMPs actively degrade nascent CaMKII*α* under stimulated conditions *in vivo*. Interestingly, inhibiting NMP activity under the basal condition also resulted in a significant increase in nascent CaMKII*α*, suggesting that NMPs degrade nascent CaMKII*α* under the basal condition, too. PSMD2 (26S proteasome non-ATPase regulatory subunit 2), which had also shown experience-driven increase in synthesis^40^, was also readily detected in the nascent protein samples. Inhibiting NMP activity significantly increased nascent PSMD2 under both basal and stimulated conditions (Fig 2C). However, no significant change in nascent PSMD2 was detected in BMI-treated samples with normal NMP function, suggesting a tight control over the level of nascent PSMD2 by NMP activity in response to increased activity. On the other hand, nascent Npas4, a well-known IEG protein, was detected at relatively low levels under basal conditions, and increased significantly with the inhibition of NMPs by BE under both basal and stimulated conditions (Fig 2C). However, no significant change in nascent NPas4 was detected in BMI-treated samples, nor was there any significant difference between BMI-BE and BMI-only samples, suggesting that NPas4 synthesis was not significantly upregulated in the tadpole brain by the BMI treatment we employed.

**Figure 3.**
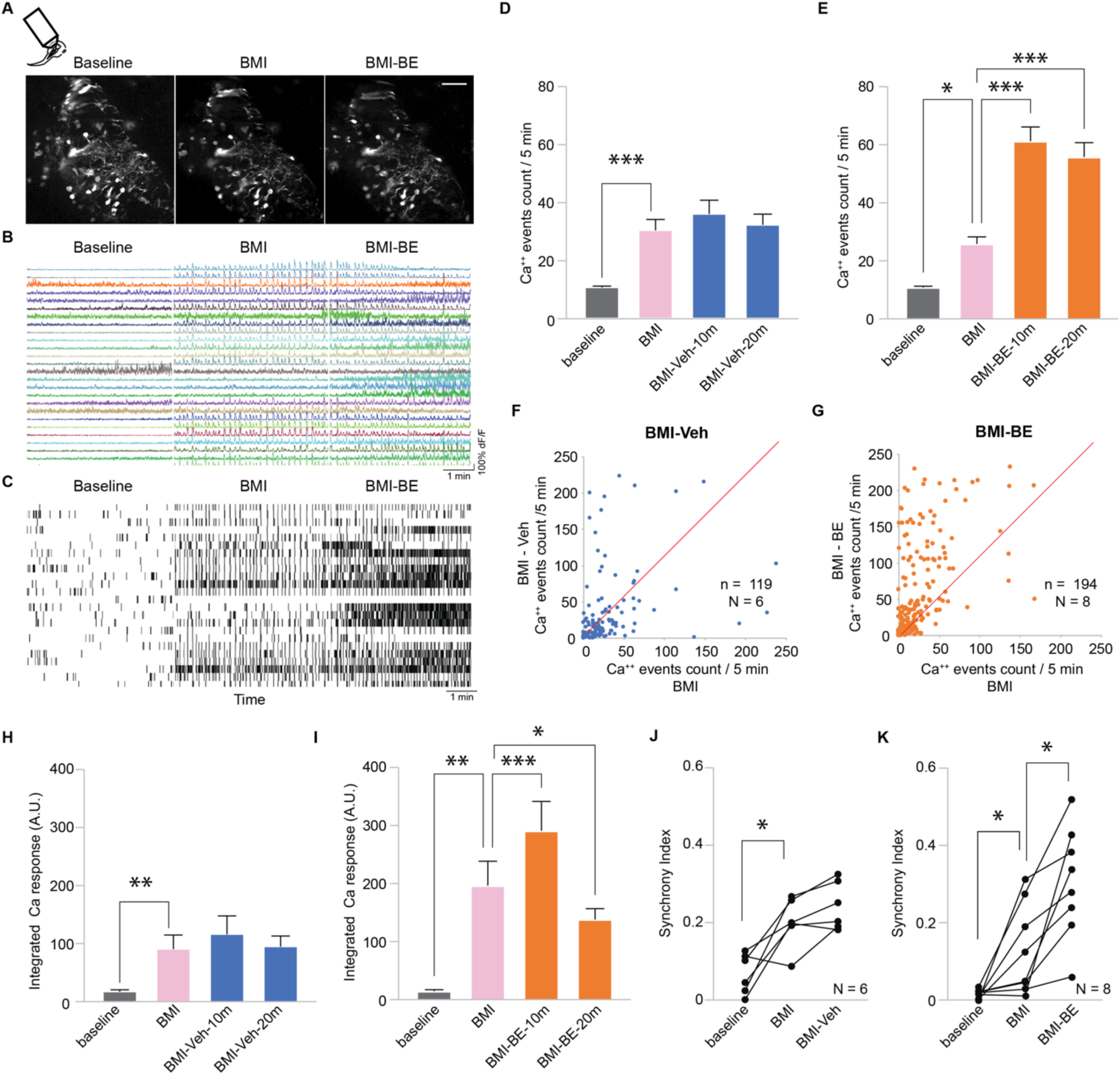
Inhibiting NMPs rapidly increases spontaneous neuronal activity and network synchrony in pharmacologically stimulated brain. A. Representative time-lapse images of GCaMP fluorescence collected in the optic tectum of a live tadpole throughout an experiment. The animal was recorded for five minutes before and after intraventricular injection of BMI, and for five minutes after BE injection. Scale bar: 50µM. B. Traces of Ca^++^ activity extracted from the GCaMP fluorescence signal in the same individual tectal neuronal soma (ROIs) during 5-min recording period following the indicated treatments. C. Raster plots (top) of Ca^++^ events derived from the data in B. D-E. Summary data of average Ca^++^ events at different time points in animals treated with BMI followed by either vehicle (D) or BE (E). F-G. Scatter plots of Ca^++^ event counts for individual ROIs over five minutes in BMI and after treatment with either vehicle (F) or BE (G). H-I. Average amplitude of integrated Ca^++^ responses in animals treated with BMI followed by either vehicle (H) or BE (I). J-K. Synchrony index calculated from Ca^++^ activity of all neurons recorded in each animal in animals treated with BMI-veh (J) or BMI-BE (K). Data points from the same animal were connected by straight lines. *: p<0.05. **: p< 0.01. ***: p<0.001. Friedman test with Dunn’s multiple comparison posthoc test. BMI-Veh: n = 119 neurons, N = 6 animals. BMI-BE: n = 194 neurons, N = 8 animals.

Notably, not all nascent proteins examined were degraded by NMPs. The newly synthesized portion of *β*-synuclein (SNCB), was not affected by the inhibition of NMP activity under either basal or stimulated condition (Fig2C), suggesting that SNCB was not degraded by NMPs in tadpole brain. No changes were detected in total protein samples of any of the proteins tested under either basal or stimulated conditions (Fig 2C), supporting that NMPs mostly degrade nascent proteins. Taken together, these data suggest that NMPs actively degrade nascent proteins *in vivo* and may have different substrate preference under different neuronal activity levels.

### NMPs actively regulate neuronal activity *in vivo*

NMPs have been shown to modulate neuronal activity in pharmacologically stimulated neuronal cultures^23^, suggesting that NMP function is involved in the regulation of neuronal activity *in vitro*. Our observation that NMPs preferentially target specific activity-regulated substrates *in vivo* also indicate a possible interaction between NMP function and neuronal activity. To determine if NMP function affects neuronal activity in intact brain circuits, we used time-lapse 2-photon Ca^++^ imaging in GCaMP6f-expressing tectal neurons to examine the effects of acute inhibition of NMPs on spontaneous neuronal activity in the brain of awake tadpoles. Ca^++^ signals recorded in neuronal soma faithfully report neuronal activity in tectal neurons^38, 50–52^. The time-lapse imaging protocol allowed us to record the activity of a population of neurons with single neuron resolution over extended periods in live animals. As shown in representative images in Figure 3A, a subset of tectal neurons can be identified at different time points throughout the time-series, making it possible to extract the Ca^++^ activity of the same individual neurons before and after each experimental treatment (Fig 3B, C).

Under the basal condition, inhibiting NMP function resulted in a dose-dependent rapid increase in spontaneous neuronal activity, as shown by the increased numbers of Ca^++^ events within the same recording time period, increased integrated signal of the compound Ca^++^ transients, as well as a moderate increase in the synchronous activity across the recorded neuronal population (Supplemental Fig 4A-D). Intraventricular injection of epoxomicin, which blocks both NMPs and intracellular 26S proteasomes, induced an increase in neuronal activity similar to that induced by BE (Supplemental Fig 4E), suggesting that the rapid increase in neuronal activity can be mostly attributed to the inhibition of NMPs.

**Figure 4.**
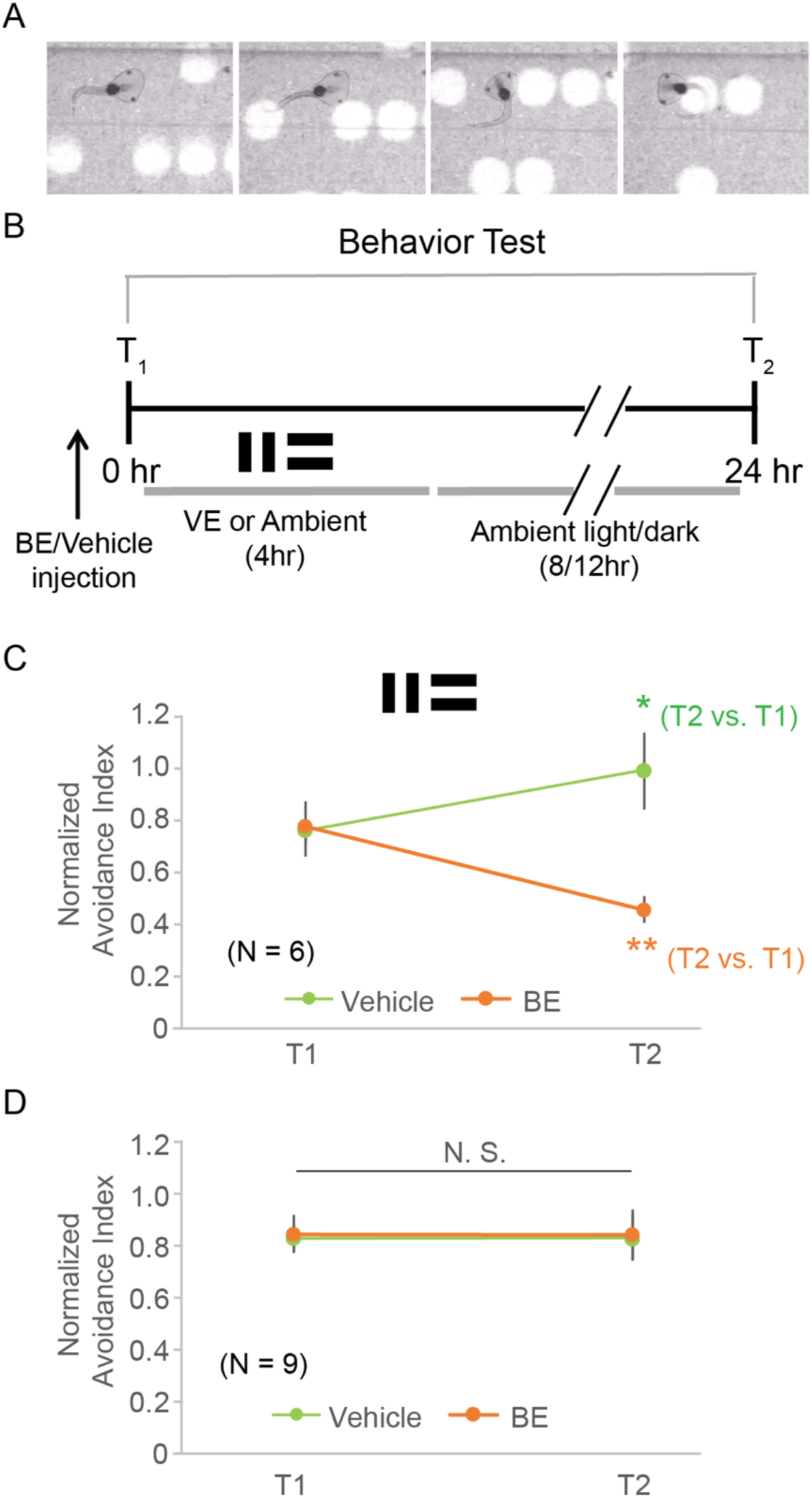
NMP activity is required for both learning-induced visuomotor behavioral plasticity and the maintenance of normal visuomotor behavior following enhanced visual training in tadpoles. A. Illustration of the visual avoidance behavior in tadpoles. Animals make a sharp turn in their swimming trajectory when they encounter an approaching visual stimulus. B. Experimental time line for visual training and behavioral test schedule. Animals first received a ventricular injection of either BE or vehicle (Veh) and were tested 30min later for baseline visual avoidance behavior performance (T1). Then animals were subjected to either 4hr of either visual training (VE) or ambient light and returned to their normal rearing conditions. Animals were tested again for avoidance behavior the next day (T2). C. Control animals improved their visual avoidance response after training (green light). Inhibiting NMPs with BE not only completely blocked learning-induced behavioral improvement following VE, but also further degraded the behavioral performance to significantly lower than pre-training baseline level. D. In the absence of VE, baseline behavioral performance was not affected by BE. *: p<0.05, **: p<0.01. Paired two-tailed Student’s t test. N: number of animal batches, marked on the bar.

As aforementioned, it is well documented that increased neuronal activity triggers an increase in protein synthesis^40, 41, 53^. As we have shown in biochemical assays, NMP activity was more prominent under stimulated conditions (Figure 2), suggesting a bigger role of NMP function in regulating neuronal activity under the stimulated condition. We therefore asked if inhibiting NMPs affects neuronal activity in the presence of BMI. As expected, blocking GABAergic inhibition in tectal neurons with intraventricular injection of BMI increased spontaneous activity levels as well as synchrony among population of neurons (Fig 3D-K). Intriguingly, inhibiting NMPs with BE following BMI treatment induced a rapid increase in neuronal activity to levels significantly above those seen with BMI alone (Fig 3E, G, I), an effect not seen in vehicle-injected animals (Fig 3D, F, H). In addition, inhibiting NMP function induced a significant increase in synchronous activity across the neuronal population on top of the BMI-effect (Fig 3 J-K), indicating a runaway of neuronal activity that could potentially disrupt activity patterns within the tectal network.

### NMP function is required for learning-induced behavioral plasticity

Two important features of the nervous system are the ability to process information and to adapt to changes in the environment, for instance through learning. Both require that neurons have the ability to detect changes in the activity pattern in response to afferent inputs above baseline neuronal activity, and the ability to maintain the neuronal activity within an intermediate dynamic range ^54–56^. Dis-regulation of neuronal activity is predicted to interfere with both normal circuit function and experience-dependent plasticity. Discoveries in multiple animal models for neurodevelopmental diseases have shown that abnormally increased neuronal activity led to hyper-synchronization and learning deficit in the neural network ^42, 43^.

Our data suggest that inhibiting NMP activity caused an abrupt increase in neuronal activity and increased synchronous activity across the network, particularly under conditions when network activity was already increased. Based on these observations, we predicted that the dysregulation of neuronal activity incurred by the inhibition of NMP function would impair experience-dependent plasticity in animals. To test this hypothesis, we used a well-established visual avoidance behavioral paradigm in tadpoles ^52, 57, 58^ to assay the effect of inhibiting NMP activity in the optic tectum on visuomotor behavior and learning-induced behavioral plasticity. The visual avoidance behavior is a tectally mediated behavior in which an animal changes swim trajectory in response to an approaching visual stimulus (Fig 4A). The percentage of avoidance events out of the first 10 encounters is quantified as the avoidance index (AI) to evaluate the behavioral performance of the animals. In addition, the performance of the visual avoidance behavior can be significantly improved in animals provided with enhanced visual training ^39, 41, 59^, likely through plasticity of the visuomotor tectal circuits ^58, 60^. This behavioral paradigm therefore can be used as a readout for both the basic circuit function and learning-induced behavioral plasticity within the visuomotor circuit. As shown in the protocol in Fig 4B, animals received a ventricular injection of either BE or vehicle and were tested shortly thereafter for their baseline avoidance performance. Animals were then subjected to either 4hr of enhanced visual training (VE) or normal ambient light, and tested again the next day to evaluate changes in their behavioral performance. Four hours of VE significantly improved the performance of avoidance behavior in animals injected with vehicle (Fig 4C), consistent with prior reports^39, 41, 59^. BE treatment completely blocked the VE-induced behavioral improvement (Fig 4C). Intriguingly, inhibiting NMPs with BE not only blocked the learning-induced behavioral enhancement, but also caused a significant deterioration of the behavior compared to the baseline level before VE (Figure 4C). In contrast, no effect on the avoidance behavior was observed in animals that did not receive VE with or without BE treatment (Fig 4D). This suggests that even though NMPs do degrade nascent proteins under basal condition, NMP function is not essential for the basal circuit function required for visual avoidance behavior. On the other hand, these results suggest that NMP-mediated degradation of activity-induced nascent proteins is essential for the expression experience-dependent behavioral plasticity and that inhibition of NMP function is particularly disruptive to circuit function under conditions with heightened neural activity.

## DISCUSSION

Proteolysis is a tightly-regulated cellular process. Although it is well documented that a well-balanced regulation of protein synthesis and degradation is required for the expression and maintenance of various forms of activity-dependent plasticity^6, 7, 32, 33, 61–64^, our understanding of mechanisms employed by neurons to achieve such a balance remains incomplete^65^. Here we report a new mechanism of proteostasis regulation employed by the vertebrate brain that controls the accumulation of activity-induced nascent proteins and may be involved in the fine tuning of proteostasis especially in response to elevated neuronal activity. We demonstrated that NMPs are present in the tadpole brain and actively degrade nascent proteins *in vivo* and found that acute inhibition of NMP activity rapidly increases neuronal activity. This suggests that the NMP-mediated degradation of nascent proteins may underlie a rapid homeostatic regulatory mechanism that helps maintain neuronal activity in check in the face of rising activity. Furthermore, we showed that inhibition of NMP function aborted learning-induced behavioral plasticity in a visual avoidance behavior paradigm in tadpoles, likely resulting from disorganized neural network activity. To our knowledge, this is the first *in vivo* demonstration of the involvement of proteolysis of nascent proteins in the regulation of neuronal activity and plasticity. New proteins are constantly made, either constitutively or triggered by changes in activity. Here we show that in addition to the well-documented UPS and autophagic pathways for protein degradation^1, 33^, and mechanisms regulating protein synthesis^66^, nascent proteins can also be immediately degraded by the ubiquitin-independent NMP pathway. This offers an additional level of regulatory vigilance over the dynamic maintenance of the fine balance needed for proteostasis in neurons, especially in face of activity fluctuations, when protein synthesis changes rapidly.

Our immunohistochemistry and EM data indicate the presence of 20S proteasomes in the neuronal membrane. The lack of biotin signals from injected BE in the neuroprogenitor cell layer along the ventricle supports the notion that NMPs are predominantly expressed in neurons. Non-neuronal cell types (such as astrocytes, microglia, etc.) as well as the processes of radial glia cells are present in the neuronal cell layers of the optic tectum^67^. Our data do not exclude the possibility that NMPs are also expressed in these other cell types, although they appear to constitute a relatively small proportion of cells in the optic tectum. In addition to their presence in neuronal somata, proteasomes have also been detected in neuronal processes, including both dendrites and axons^31, 68–70^. The ultrastructural localization on NMPs at tectal synaptic sites, similar to what has been reported in mouse hippocampal neurons^23^, suggests that NMPs are positioned to be directly involved in the timely degradation of locally translated activity-induced proteins, which can be particularly important due to the limited volume capacity and the immediate spatial proximity to the site of the action with regard to synaptic transmission^71^. It has been proposed that synaptic proteins are made in excess but those that are not incoporated into synapses are readily degraded^72^. The localization of NMPs at the perisynaptic plasma membrane supports a particularly attractive mechanism for such rapid timely degradation.

Increased neuronal activity is well known to induce significant increases in protein synthesis in neurons^53, 65, 71^. If NMPs function to degrade proteins produced in excess, one might expect that the total amount of nascent proteins degraded by NMPs would be higher under stimulated conditions than that under the basal condition. Indeed, we observed that the NMP-mediated nascent protein degradation is more prominent with increased neuronal activity. Estimated by the dot blot data, the percentage of total nascent proteins degraded by NMPs was ∼15% (Fig2B, BMI-BE subtract BMI) of the control total nascent protein population under the stimulated condition and was undetectable under the basal condition. This represented about a third of the total amount of nascent proteins induced by the increased neuronal activity (∼43% of control, BMI-BE subtracted by control). In addition, when individual proteins were examined, the percentage degraded by NMPs could be much greater (Figure 2C), suggesting that NMP-mediated degradation of nascent proteins is not ubiquitous, but instead may function to degrade excessive nascent protein at a speed in accordance with the production rate of individual nascent proteins to maintain the level of each individual protein at its physiological range.

The observation that different individual nascent proteins may undergo NMP-mediated degradation at different levels under basal and stimulated conditions is very interesting. This indicates that NMPs may differentially degrade nascent proteins under different activity levels on a substrate-specific basis. The individual proteins we examined were selected based on prior reports of their involvement in activity-dependent protein synthesis. CaMKII*α* is a key molecule underlying synaptic plasticity^73^. It is well-documented that both the mRNA and protein levels of CaMKII*α* are upregulated in neurons in response to activity^6, 40, 41, 74–76^. CaMKII*α* mRNA is localized to dendrites, and is also rapidly recruited to dendritic spines and translated into CaMKII*α* protein when activity increases^76–79^, making it a plausible candidate substrate for NMPs. Indeed, our data clearly demonstrate that CaMKII*α* was synthesized under basal resting condition and its synthesis increased significantly within 1hr of pharmacological stimulation. And the nascent CaMKII*α* significantly increased when NMPs were inhibited under both resting and stimulated conditions, suggesting that CaMKII*α*is a substrate of NMPs. The proteasome subunit PSMD2 had been reported to show upregulated synthesis in tadpole brain following 4hr of enhanced visual stimulation^40^. The synthesis and degradation of proteasome components are known to be regulated by neuronal activity in the brain^21^. In the one-hour pharmacological stimulation paradigm we employed here, with normal NMP function, no change in nascent PSMD2 was seen in the stimulated samples compared to the basal condition. However, inhibiting NMPs significantly increased the amount of nascent PSMD2, suggesting that NMPs actively degraded the newly-synthesized PSMD2 to maintain its basal level. Npas4 is a well-known protein product of an IEG that is transcribed and translated in response to upregulated neuronal activity^22, 80–82^. Under basal conditions, nascent Npas4 was detected at relatively low levels. However, when NMP was inhibited, we observed a significant increase in nascent NPas4. This suggests that Npas4 is constitutively synthesized, but NMP-mediated rapid degradation helps maintain the nascent protein at a low level. Maintaining low levels of constitutive expression of IEG proteins is essential to ensure a rapid response to salient stimulus^82–85^. We did not observe any significant increase in the nascent NPas4 protein level under the stimulation protocol we employed, suggesting that one hour treatment of BMI was not an optimal stimulation in tadpole brain to induce a significant increase in Npas4 synthesis. Taken together, these data suggest that NMPs may function as a mechanism for rapid (possibly instantaneous) control of the nascent protein production. And the NMP-mediated degradation of individual proteins varies under different activity levels in a protein-by-protein manner. Whether such differential NMP-mediated degradation of different nascent proteins is due to differences in the substrate selectivity of NMPs or differences in the levels of protein synthesis (and therefore the availability of nascent protein substrate) of individual proteins remains to be elucidated.

Our results demonstrated that inhibiting NMP function caused rapid dysregulation of neuronal activity that led to hyper-synchronization of the neuronal population, and abolished visual training-induced behavioral plasticity in the tadpoles. The epoxomicin binding to the 20S core particle is irreversible^47^ and our biochemistry data showed that BE remained bound to the NMP for at least 6hr after injection (Fig 1C), ensuring that the NMPs remain blocked during the VE training period. This suggests that enhanced visual training, while enabling the visuomotor circuit to be refined and strengthened for better behavioral performance, may also render the circuit labile to perturbations. The formation of new memories is thought to be the result of a series of fine-tuned modifications that occur in a highly specific manner at various synapses involved in the bebavior, from sensory inputs to motor output^86^. The training process triggers strengthening of some synapses and weakening of others, thus changing the detection threshold to certain sensory inputs and eventually the response manifested by animal behavior. Mechanisms of synaptic plasticity have been shown to trigger such plastic changes at the synapses. Interestingly, proteasome inhibition has been shown to enhance the induction of LTP but blocked expression of late phase LTP^61^. Even though this effect was attributed to the UPS pathway, it is noteworthy that the proteasome inhibitor used in the study, epoxomicin, also inhibits NMPs. As we have shown here, application of epxomicin induced a rapid increase in spontaneous neuronal activity that is very similar to what we observed with the application of the specific NMP inhibitor, Biotin-Epoxomicin (supplemental figure). The increased activity induced by acute inhibition of NMPs may lead to a transient further increase in activity that could be interpreted as enhanced LTP induction. What’s more, the effect of proteasome inhibition on late phase LTP was blocked by the protein synthesis inhibitor anisomycin^61^, indicating the underlying mechanism involves newly synthesized proteins. Similarly, blocking protein synthesis in tadpole brain also ablated the learning-induced visuomotor behavioral plasticity^41^ studied here.

What might be the mechanism underlying the role NMPs play in regulating neuronal activity? Runaway neuronal activity disrupts the activity patterns in neural networks that are essential to form synaptic-specific modifications of neuronal connectivity, which is the basis for learning and memory^87^. Maintaining neuronal activity in an intermediate range is the core function of homeostatic plasticity mechanisms. Activity-induced nascent proteins trigger subsequent signaling pathways for the expression of synaptic plasticity^82, 84, 88^. Intuitively, timely degradation of these nascent proteins may hold such activity-induced synaptic changes in check, preventing them from plunging the network into the destabilizing positive-feedback loops. NMP-mediated degradation of activity-induced nascent proteins may underlie a fast homeostatic regulation of neuronal activity, particularly in face of elevated activity levels. Activation of CaMKII following increased synaptic activity leads to phosphorylation of serine 120 which in turn leads to enhanced 26S proteasome activity^89^, generating a potentially positive feedback loop within the signaling cascade. Putting a cap on the amount of newly synthesized CaMKII will help to put a brake on the downstream signaling cascade. The activity-dependent regulation of proteostasis in neurons is likely a highly intricate and delicate process. Activity has been shown to translocate proteasomes to active dendrites in hippocampal neuronal cultures^2^. Increased levels or function of certain plasticity-related proteins may involve activity-induced degradation of negative regulator. For example, NMDAR activation in neurons has been shown to induce proteasomal degradation of MOV10, a synaptic translational repressor, which in turn led to increased expression of CaMKII*α*^75^. Likewise, tight control of the newly-synthesized proteasome subunits such as PSMD2 could be part of the dynamic regulatory scheme underlying activity-dependent proteostasis. Failure to remove the excessive amount of such activity-induced nascent proteins may not only jeopardize the expression of synaptic plasticity but also lead to damage in the existing circuitry and impair normal information processing, as shown by the detrimental effect of blocking NMPs on visual avoidance behavior following enhanced visual training we observed.

Another potential pathway through which NMPs function can affect neuronal activity is by the peptides generated by the 20S catalytic degradation of nascent protein substrates^1, 23^. Our observation that inhibiting either protein synthesis or degradation *in vivo* both resulted in a transient increase in neuronal activity suggests that there might exist some common factors regulating neuronal activity that are generated by both protein synthesis and degradation. The peptides produced by the degradation of nascent protein can be a reasonable candidate for such factors, although currently very little is known about the identities and functional property of these peptides. It is possible that multiple mechanisms underlie the functional involvement of NMPs in regulating neuronal activity. More data are emerging suggesting different activity patterns can induce different compositions of the nascent transcriptome and proteome, as a result of the recruitment of different classes of neurons, involvement of a differential neural circuitry, and even induction of different plasticity mechanisms, both *in vivo* and *in vitro*^9, 40, 90, 91^. The functional contribution of NMP-mediated degradation of activity-induced nascent protein may also vary under different activity paradigms depending on the nascent proteome induced.

## MATERIAL AND METHODS

### Key resources table

| Reagent type | Description | Source or reference | Identifiers |
| --- | --- | --- | --- |
| Antibody | Mouse-anti-CaMKIIα | Novus | Cat#NB100-1983,<br>RRID:AB_10001339 |
| Antibody | Rabbit Anti-b-Synuclein | Sigma-Aldrich, | Cat# SAB4502827,<br>RRID:AB_10761178 |
| Antibody | Anti-Actin | Sigma-Aldrich<br>(Millipore) | Cat#MAB1501,<br>RRID:AB_2223041 |
| Antibody | Mouse Anti-B-Tubulin | Sigma-Aldrich | Cat#T8535,<br>RRID:AB_261795 |
| Antibody | Rabbit Anti-PSMD2 | Cell Signaling<br>Technology | Cat#25430S,<br>RRID:AB_2798903 |
| Antibody | Rabbit Anti-BSA | Invitrogen | Cat#PA1-29262,<br>RRID:AB_1956427 |
| Antibody | Goat anti-Biotin | Fisher Scientific | Cat#31852,<br>RRID:AB_228243 |
| Antibody | Rabbit anti-NPAS4 | Gift from<br>Dr. Yixing Lin |  |
| Antibody | Mouse anti-Ubiquitin | Fisher Scientific | Cat#131600,<br>RRID:AB_2533002 |
| Antibody | Goat Anti-Rabbit IgG<br>(H+L) HRP | Thermo<br>Scientific | Cat#31466,<br>RRID:AB_10960844 |
| Antibody | Goat Anti-Mouse IgG<br>(H+L) HRP | Thermo<br>Scientific | Cat#31431,<br>RRID:AB_10960845 |
| Antibody | Rabbit Anti-Goat IgG<br>(H+L) HRP | Thermo<br>Scientific | Cat#A16136,<br>RRID:AB_2534807 |
| Antagonist | Biotin-Epoxomicin | Ramachandran<br>et al., 2017<sup>23</sup> |  |
| Antagonist | Bicuculline Methiodide | R&D systems | Cat# 2503 |
| Antagonist | Epoxomicin | Apexbio | Cat# 50-190-4754 |
| Antagonist | Anisomycin | Sigma-Aldrich | Cat#A5862 |

### Animals

Albino Xenopus laevis embryos were obtained from either in-house mating or from Xen Express (Brooksville, FL), and reared at 21-22°C with 12 hr dark/12 hr light cycle in 0.1×Steinberg’s solution (in mM: 58.0 NaCl, 0.67 KCl, 0.34 Ca(NO_3_)_2_, 0.83 MgSO_4_, 3.0 HEPES, pH 7.2). Animals were fed from stage 47 (Nieuwkoop and Faber, 1994). All animal protocols were approved by the Institutional Animal Care and Use Committee (IACUC) of the Scripps Research Institute and Georgetown University IACUC. Stage 47-48 tadpoles of either sex were used for all experiments.

### Intraventricular injection

For intraventricular injection of different drugs (BMI, BE, Epox, Ani) or the non-canonical amino acid AHA, animals were anesthetized with 0.02% MS-222 (Sigma-Aldrich, Cat#A5040) and placed on a moist Kimwipe under a dissecting microscope. The corresponding drugs or vehicle solution (as indicated in each experimental condition) was pressure-injected through a glass micropipette into the midbrain ventricle using a Picospritzer. The micropipettes were calibrated (Sive et al., 2010) to inject 20-25nl of solutions each time. The total ventricle volume of stage 47-48 Xenopus laevis tadpoles is estimated to be about 200∼300nl on average, inferred from a prior study (Mogi et al., 2012). The concentration of solutions injected was listed as the injection concentration, not the final concentration *in vivo*, due to difficulties in precisely assessing the total final volume (i.e. total ventricle volume plus the interstitial volume of the brain tissue).

### Immunohistochemistry

Tadpoles which received intraventricular injections of BE were fixed 1hr post-injection for immunocytochemistry together with un-injected controls. To maximize the direct detection of injected BE in the tadpole brain, high concentration (1mM) of BE was injected for ISH and membrane preparation experiments described in figure 1. Lower concentrations of BE (5-250µM as indicated for each experiment respectively) were injected for subsequent biochemical, functional imaging, and behavioral experiments. Animals were fixed with 4% paraformaldehyde freshly diluted from 16% stock solution (Electron Microscopy Sciences, Fort Washington, PA) in 1×PBS (pH 7.4), using a Pelco BioWave Pro microwave (Model 36500, Ted Pella, Redding, CA. 350 mV on 20 sec, off 20 sec, on 20 sec, followed by 150 mV on 1 min, off 1 min, on 1 min). The animals were then post-fixed at 4°C overnight and washed in 1×PBS using the microwave (150 mV on-off-on, 1min each). 30 µm vibratome sections of the dissected brains were cut for free floating immunofluorescence labeling. Sections were incubated in 1% Sodium Borohydride (Sigma) in 1×PBS for 15 min to quench auto fluorescence, blocked in 10% normal goat serum (Jackson Lab, ME) and 1% of fish gelatin (Bioworld) in PBS with 0.1% tween-20 (PBST) for one hour in room temperature, followed by incubation in goat anti biotin polyclonal antibody (Invitrogen, Cat # 31852, RRID: AB_228243, 1:500 in PBST with 1% normal goat serum and 1% fish gelatin) for 72 hours at 4°C. Secondary antibody (donkey anti goat Alexa Fluor 488, Thermo Fisher Scientific Cat# A-32814, RRID: AB_2762838) was diluted 1:500 in PBST with 1% normal goat serum and 1% fish gelatin and incubated overnight at 4°C. After 3×15 min rinses with PBS, sections were mounted on slides in Vectashield Mounting Medium with DAPI (Vector Laboratories, Burlingame, CA). Brains from BE-injected and un-injected groups were embedded in the same blocks and processed under exactly same conditions throughout the experiments. Images of immunolabeled sections were collected on a Nikon C2 confocal microscope with a 40× PlanFluor Oil objective (N.A. 1.3) under exactly same acquisition parameters.

### Electron Microscopy

Stage 47 tadpoles were anesthetized by immersion in 0.02% MS-222 and then a fixative consisting of 3.5% paraformaldehyde, 1% glutaraldehyde and 0.02% CaCl_2_ in 0.035M sodium cacodylate buffer was pressure injected into the ventricle. Then, tadpoles were immersed in fixative and fixed using two bouts of microwave fixation at 150W for 1min, followed by overnight fixation at 4°C. Tadpole brains were dissected, embedded in an albumin-gelatin mixture, and sectioned at 50m on a vibratome. Sections were blocked and permeabilized in 5% normal donkey serum, 1% fish gelatin, and 0.1% Tween-20 for 1h at room temperature. Then, sections were incubated in 1:500 mouse anti-proteasome 20S α1,2,3,5,6&7 subunits (Enzo, BML-PW8195, RRID: AB_10541045) for 3 days at 4°C followed by 2h in 1:800 anti-mouse FluoroNanogold (Life Technologies, A24920) at room temperature. GoldEnhance EM Plus (Nanoprobes, 2114) was used to enhance Nanogold particle size for visualization. The enhancement duration was calibrated so that primary and no primary antibody groups contained gold particles of similar size after enhancement.

Sections were post-fixed in 1% osmium tetroxide for 1h, dehydrated in acetone series (50%, 70% with 4% uranyl acetate, 90%, and 100%), and infiltrated with EMbed-812 resin (50% in acetone and 100%) (Electron Microscopy Sciences, 14120). The next day, sections were flat-embedded in 100% EMbed-812 resin between two sheets of Aclar plastic (Electron Microscopy Sciences, 50425) and incubated at 65°C overnight. Ultrathin sections (60nm) were cut using a diamond knife and collected on formvar-coated nickel slot grids (Electron Microscopy Sciences, FF2010-NI). Ultrathin sections were examined with a FEI Talos L120C electron microscope and photographs were taken with a Ceta 16M CMOS camera at 8,500X or 22,000X magnification. In electron micrographs, membranes were pseudocolored blue using Photoshop (Adobe) for easier visualization.

### Membrane preparation and western blotting

Tadpoles (40 in each group) received intraventricular injections of 1mM BE or 1mM epoxomicin (Epox) and brains were dissected either 30 minutes or 6 hours later. Membrane preparation was performed for biochemical analysis as previously described^23^. In brief, frozen tadpole brains were homogenized in hypotonic lysis buffer (5 mM HEPES, 2 mM ATP, 1 mM MgCl2). Nuclei were pelleted at 800 r.p.m. for 5 min, and the supernatant containing membranes was pelleted at 55,000 r.p.m. for 1 h. Pelleted membranes were washed twice and re-pelleted for the final membrane fractionation sample. Supernatants containing the cytosolic extracts were concentrated down to the same volume that membranes were eventually resuspended in. Both membrane and cytosolic extracts were boiled in Laemmli buffer for 10 min. Equal volume of both samples were loaded on to SDS-PAGE gel and processed for western blotting.

### BONCAT and immunoblotting

#### Sample processing and data analysis for Dot-Blot

1M Azidohomoalanine (AHA) stock solution was made from powder (Click Chemistry Tools, Cat# 1066100) in 1xPBS with pH adjusted to 7 using 1N NaOH. All drug solutions were prepared to have a final concentration of 350mM AHA and contain ∼0.01% fast green dye for injection visualization. Epoxomicin (Epox, Apexbio, Cat# 50-190-4754, 50∼100µM), Biotin-Epoxomicin (BE, synthesized de novo and purchased from the Leiden University Institute of Chemistry with purity and verified as described before (Ramachandran et al., 2017), 50∼100μM), Anisomycin (Sigma-Aldrich, Cat#A5862, 25 μM) and Bicuculline Methiodide (BMI, R&D systems, Cat# 2503, 25∼50μM) were injected into the midbrain ventricle of st48 tadpoles. 30 minutes after injection, the brains were dissected. 22 animal brains were collected for each experimental group and stored at -80°C to be processed the following day. Tadpole brain tissue were processed as previously described (Liu et al., 2018). In brief, brains homogenized in 1xPBS containing 1% SDS and protease inhibitors (Complete EDTA-free Protease Inhibitor cocktail, Roche, Sigma Aldrich, Cat#11873580001) and boiled for five min, followed by sonication for 30s. Next, protein concentration of the lysates was measured using the BCA Protein Assay Kit (Thermo Fisher Scientific, Cat#23227) and the samples diluted to 1mg/mL. A small portion of the lysate was boiled for 10 min with 2X SDS Laemmli sample buffer (Bio-Rad, 1610737) with BME and saved as total protein samples. For a 100 μl of click chemistry reaction, 86.5μL of the lysate was incubated with 13.5 μl of click mix solution (1.7mM Triazole ligand in 4:1 tBuOH/DMSO (Sigma), 50 mM CuSO4 (Sigma), 5 mM Biotin-PEG4-Alkyne (Click Chemistry Tools, Cat#TA105100) and 50 mM TCEP (Sigma) added in that order). The reaction was allowed to proceed for 90 minutes at room temperature. Then 50μL of 9M Urea (Millipore Sigma, Cat#66612-M)/75mM Ammonium bicarbonate (Millipore Sigma, Cat#A6141) and 150 μL of 2X Laemmli sample buffer with BME were added to the click reaction mix and incubated for 30 min at RT. For dot-blot, protein samples were serially diluted with 1xTBS (1:2, 1:4 & 1:8) and loaded in triplicates onto a nitrocellulose membrane assembled in a 96-well dot-blot apparatus (Biorad, Cat#1703938). The membrane was then stained with Ponceau (Sigma-Aldrich, Cat# 09189) and imaged. Then the membrane was blocked in 5% non-fat dry milk (Bio-Rad, 1706404) in 0.1%TBS-Tween for one hour and incubated with anti-Biotin (Invitrogen, Cat # 31852, RRID: AB_228243, 1:1000) overnight at 4°C, followed by 1hr of secondary antibody incubation at room temperature and 3X washes with 0.1% TBST. The blot was then developed with Clarity Western ECL Substrate (Cat# 1705061). Images were acquired with ImageQuant LAS 4000 (Amersham, GE-Healthcare Life Sciences) and densitometric analysis was done on Image J (NIH) using the Dot-Blot Analyzer. The biotin signal of each dot was first normalized to its corresponding ponceau signal for loading control and then normalized to the control sample from the same batch of experiment. Multiple exposures of each blot were obtained to ensure that the quantified blots were within the linear range.

#### Purification of biotinylated AHA-tagged nascent proteins and data analysis for Western Blot

Tadpole brain lysate and click chemistry were performed in the same way as described for dot-blot sample processing. To serve as an internal loading control, 1.5μL of 0.1mg/mL biotinylated BSA (BioVision, Cat#7097-25) was added to each tube of click mix. Once the click reaction was complete, proteins were purified by methanol/chloroform/water precipitation. The protein pellets were air dried and resuspended in 100 μL of 6M Urea/25 mM ammonium bicarbonate/0.5% SDS in PBS and vortexed for 20-30 min till pellets completely dissolved. 25μl of pre-washed Neutravidin beads slurry (Pierce High Capacity Neutravidin Agarose Resin ThermoFisherScientific, Cat#29202) were added to each protein sample and final volume was brought to 1000μl with PBS. The tubes were rotated head-over-head overnight at 4°C. The next day, the Neutravidin beads were washed with PBS, then incubated in 1% SDS in PBS for 10 min, followed by two more washes with PBS. Following the final wash, the beads were boiled in 30 μL of 2X Laemmli sample buffer w/BME for 15 min. The solution was then centrifuged at 14,000 rpm for two min and the supernatant was collected as the purified biotinylated nascent protein samples. The nascent and total protein samples were then run on a 4-12% gradient SDS-polyacrylamide gel along with Precision Plus Protein Kaleidoscope ladder (Bio-Rad, Cat #1610395). Then the proteins were transferred onto nitrocellulose membranes using either a Bio-Rad Trans-Blot Turbo Transfer system (25V, 2.5 A, 9 min) or an Invitrogen iBlot 2 Dry Blotting system (P0 program modified as 20V for 1 min, 23V for 4 min and 25V for 4 min). Once transfer was complete, the membrane was cut horizontally into three strips right above the 75kDa and 37kDa ladder standards and each strip was incubated with the appropriate primary antibody solution overnight after 1hr of blocking in 5% milk in 0.1%TBST. Then the strips were washed 3X with 0.1% TBST and incubated with the corresponding secondary antibodies. Clarity Western ECL Substrate (Cat# 1705061) was used for blot development and images were acquired with ImageQuant LAS 4000. Protein band quantification was performed by densitometry analysis using ImageJ (NIH). Multiple exposures were taken to ensure the quantified bands were within the linear range. For total protein samples, the band intensity was first normalized to tubulin signal from the same sample for loading control and then normalized to the value of the control sample from the same batch. For biotinylated nascent protein samples, the band intensity was first normalized to the BSA loading control that was added at the time of the click reaction, and then normalized to the value of the control sample from the same batch. Certain immunoblots were stripped with Restore Plus Western Blot Stripping Buffer (Thermo Scientific, 46430) for subsequent rounds of antibody probing. For neutravidin pull down of the BE-injected samples with epoxomicin pre-injection (no AHA labeling), neutravidin beads were directly added to the total lysate of brain tissues and processed the same as described above.

#### Antibodies

Antibody dilutions were made according to manufacturer’s directions unless otherwise specified. See key resource table for details.

### *In Vivo* Two-Photon Imaging of Spontaneous Ca++ Activities in Tectal Neurons and Data Analysis

For functional imaging of spontaneous calcium activity, we co-electroporated animals at stage 46-47 with pGP-CMV-GCaMP6f (2 mg/mL, Addgene plasmid # 40755) and CMV-turboRFP (1mg/ml). Electrical current was delivered through a Grass SD9 stimulator (34V, 1.6 msec duration), 3-5 pulses were applied with each polarity. Animals were pre-screened 3-5 days after electroporation for those with moderate to high number of transfected cells expressing turboRFP. On the day of imaging, the animal was immobilized with pancuronium dibromide (1mM in 0.1X Steinberg solution; Apexbio, Cat#15500-66-0) for 1 minute^92^. The animal was then placed in a customized 35mm petri dish with sylgard-carved imaging chamber and covered with 1% low-melting agarose (Lonza #50101, Rockland, ME) for extra immobilization. The petri dish was then filled with freshly aerated Steinberg’s solution during Ca^++^ imaging. Time series of spontaneous Ca^++^ activity was collected at 30Hz frame rate either on a Scientifica multiphoton resonant microscope (Scientifica, UK) with a 25x water immersion objective (Olympus ultra 25xMPE, 1.05NA) or a Bruker Ultima Investigator multiphoton microscope (Bruker, Billerica, MA) in resonant scanning mode with a 20x water immersion objective (Olympus XLUMPLFLN20XW, 1.0NA). Wavelength of 940nm was used to excite GCaMP6f. The optic tectum was imaged with a zoom factor of 2. At each time point, animals were imaged for five min in the dark. For each experiment, a five-minute baseline was taken before the animal received intraventricular injections for subsequent treatments. Animals were kept in the agarose throughout the whole experiment to ensure that the same optical plane (with the same population of GCaMP6f-expressing neurons) could be imaged for all time points. The patterns of turboRFP-expressing cells were used as landmarks to help identifying the imaged optical plane.

Time-series of calcium response data were processed in Image J (NIH) by manually identifying the region of interest (cell soma) and then further analyzed with customized Matlab scripts. Whenever needed, the original image stacks were aligned using the registration function of the calcium imaging toolbox suite2p (RRID:SCR_016434)^93^ before the analysis in Imag J. Only neuronal somas, which were identified by their anatomical location and cell morphology, were included in the data set. The extracted GCaMP6 fluorescence was used to calculate dF/F based on an exponentially weighted moving average algorithm^94^ to remove the slow drifting of baseline signal as well as fast oscillatory noises resulted from occasional tissue pulsation. Spontaneous Ca^++^ event was defined as 3SD above the mean dF/F in the GCaMP fluorescence during the five-minute baseline recording session immediately before the first drug injection^95^. The same threshold (mean dF/F + 3SD during the baseline session) was applied to dF/F data of all subsequent time points recorded from the same neuron. The total number of spontaneous Ca^++^ event peaks during each 5-min recording session were taken as Ca^++^ event count. The integrated Ca^++^ response was calculated as the integrated area under the curve above threshold for each Ca^++^ event. Synchronous Ca^++^ events were analyzed based on previously published method^42, 96^. In brief, the denoised dF/F trace of the whole time series were binarized by the threshold, i.e. everything above the threshold was set to 1 and the rest was set to 0. The binarized Ca^++^ traces of all neurons recorded in the same animal were then summed up to represent the Ca^++^ activity in the population. A permutation of 1000 trials was then performed by circularly shuffling the binarized Ca^++^ trace of each neuron, which randomize the timing of all Ca^++^ events without changing the total activity level of the neuron. The 95^th^ percentile value of all summed population Ca^++^ traces was then defined to be the threshold for synchrony. The synchrony index for each animal at each time point was calculated by the average of all events from all ROIs that were above the synchronous threshold. The average was taken so that the value can be compared across different animals with various numbers of ROIs recorded. The synchronous index was a value between 0 and 1, with 0 being no synchrony, and 1 being all cells were synchronously active. For example, a sync index value of 0.2 means 20% of the whole time period there were synchronous events going on in the neuronal population.

For BMI-BE data presented in figure 3, treatments with 50∼250 μM BE showed a consistently rapid increase in neuronal activity on top of BMI-induced heightened neuronal activity across animals from different clutches, thus we pooled data from all animals injected with 50∼250 μM BE following the BMI-treatment for the BMI-BE dataset.

### Visual Avoidance Assay and Visual Training

The visual avoidance assay was conducted as previously described^57^. In brief, Stage 47 tadpoles were prescreened a day before to ensure they display normal visuomotor response and only animals passed the screening were included in the following experiment. On the day of the experiment, tadpoles were placed in an 8×3 cm testing chamber molded with multiple clear lanes filled with rearing solution to a depth of ∼1cm, with one tadpole in each lane. A random pattern of moving dots of 0.4cm diameter was projected onto a back-projection screen on the bottom of the testing chamber with a microprojector (3M, MPro110). Tadpoles were visualized with infrared LEDs and videos of tadpole behavior were captured with a Hamamatsu ORCA-ER digital camera^97^. Visual stimuli were generated and presented by MATLAB. To quantify the visual avoidance responses, videos were manually analyzed post hoc. An avoidance response was scored when a tadpole displayed a sharp turn within 500 ms of a dot moving perpendicularly toward the eye. The Avoidance Index was quantified as the fraction of avoidance responses out of the first ten total encounters. Tadpoles were only given a score if they swam normally and if they had 10 or more moving dot encounters during the testing period. Tadpoles were then evenly split into experimental and control groups based on their Avoidance Index score such that the average baseline score for each experimental group was the same. Animals were assigned starting with the highest scoring and working towards the lowest until the desired number of animals for each group was reached. Animals with low scores or that did not reach ten events were therefore less likely to be assigned. The lowest scoring (and un-scorable) 50% of the tested tadpoles were not assigned to groups and excluded from the experiment.

Following group assignment, tadpoles were left to rest for approximately three hours before getting intraventricular injection with either vehicle or BE (50mM). The first behavior timepoint (T1) was taken 20-40 minutes after the injection to allow animals to completely recover from the anesthesia. Immediately after T1, the group assigned for enhanced visual training (VE) were exposed to visual stimulus training with random drifting gratings (1 cm width; 0.3 Hz; Luminance: 25 cd m^-2^) at all four directions for four hours^39, 41, 59, 98^. The control groups were left on the bench top exposed to normal ambient light for the same period of time. Then both control and VE groups were returned to the incubator with 12hr/12hr L/D lighting schedule for the rest of the day till next morning. A second timepoint for behavioral performance was taken for all animals the following morning. All data were analyzed post-hoc as described above, blind to the experimental groups.

All subsequent behavior tests at T1 and T2 following baseline were collected with two tadpoles per lane, and animals were no longer individually tracked. All other aspects of the data collection, including the stimulus and lane chamber used, remained the same as the baseline test. The effect of visual training on the improvement of visual avoidance behavior varies from batch to batch. To specifically examine the effect of NMP inhibition on learning-induced behavioral plasticity, only batches exhibited significant improvement in their avoidance behavior following visual training were included in the final dataset. The N value of the final dataset reflects the number of batches of animals.

### Statistical Test

Statistical tests were performed using GraphPad Prism version 9.2.0 for Mac OS (GraphPad Software, San Diego, California USA). Kolmogorov-Smirnov test was used to determine if the data set was normally distributed and parametric (for normally-distributed dataset) or non-parametric tests were used accordingly. All data are presented as mean ± s.e.m. Data are considered significantly different when p values are less than 0.05. Posthoc power analysis was performed using JMP Pro 15 (SAS institute). The statistical test used for each experiment is specified in the results. Experiments and analysis were performed with the experimenter blinded to the experimental conditions.

## ACKNOWLEDGEMENTS

We thank the Microscopy Core at Scripps Research Institute for their technical support. We thank Dr. Han-Hsuan Liu and Eric Carlson for their help in the initial optimization of the *in vivo* BONCAT labeling and sample processing protocol. We thank Dr. Yingxi Lin for generously sharing the Npas4 antibody and Dr. Ju Lu for generously sharing the Matlab script for synchrony analysis. This work was funded by the National Institutes of Health (KL2TR001432 to H.H; EY11261,1EY031597, NS114975 to H.T.C.; R01 NS110754 to S.S.M.)

## AUTHOR CONTRIBUTION

H.H., K.V.R., S.S.M. and H.T.C. conceived the project and designed experiments. H.H., K.V.R., N.M., R.F., A.A. and R.B. conducted experiments and analyzed the data. H.H., K.V.R., S.S.M. and H.T.C. interpreted the data and wrote the paper. All authors reviewed the final manuscript.

---

**Supplemental Figure 1.**
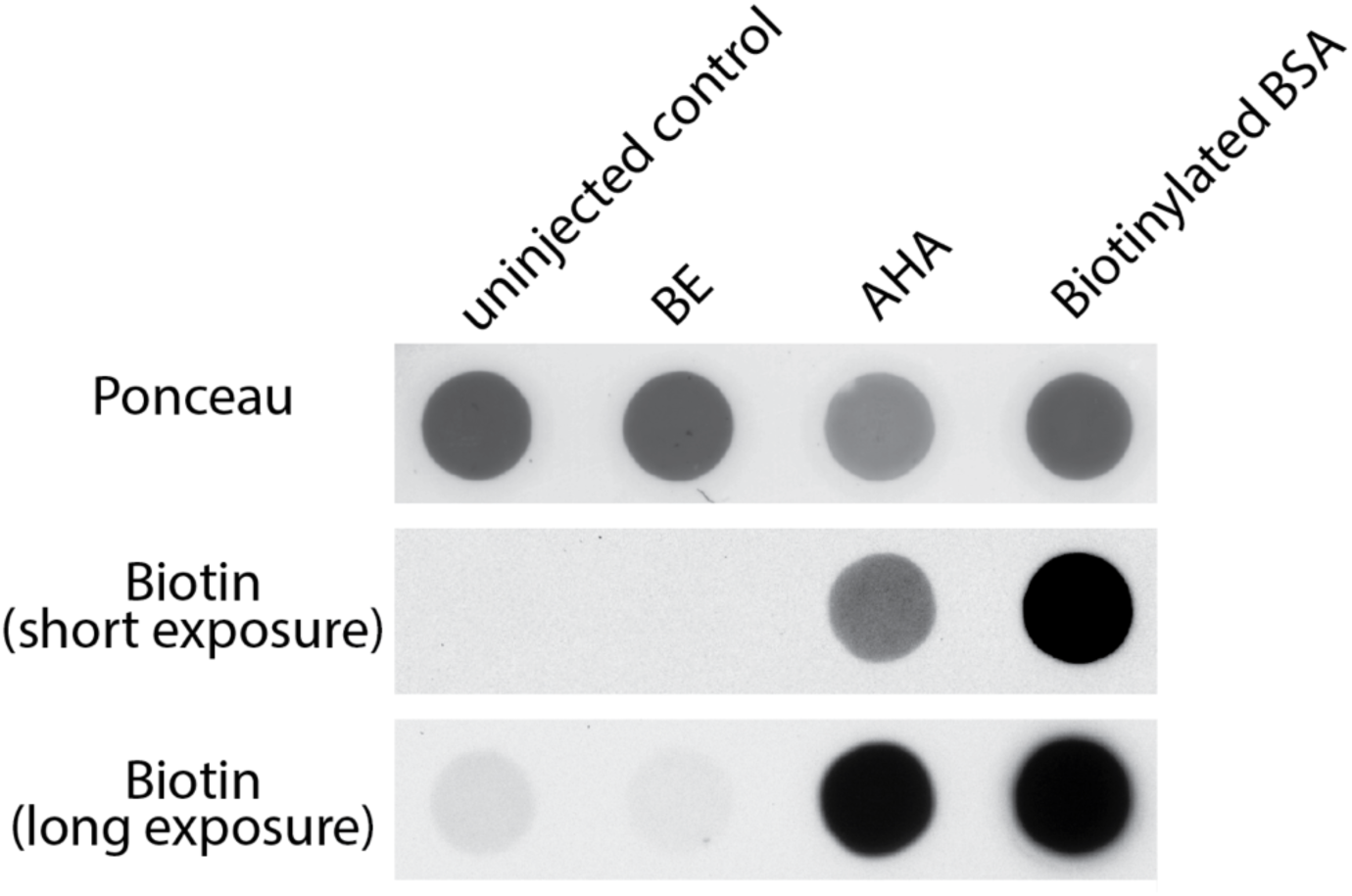
Injected BE and endogenous biotin do not interfere with the detection of AHA-biotin labeled nascent proteins by biotin antibodies in dot-blot. Protein samples from un-injected control and BE (100µM)-injected (no AHA) animals were loaded on the same blot together with samples from AHA-injected animals to blot for biotin signal. Pure BSA sample spiked with 0.3 μg biotinylated BSA was loaded as positive control. Ponceau staining show the amount of total protein loaded. No biotin signal was detected in either the un-injected or the BE-injected samples within the linear range of exposure period for the AHA-injected samples (short exposure). Even under extra-long time of exposure, when both the AHA-injected and the biotinylated BSA samples were over saturated, the biotin signals in the un-injected and BE-injected samples were negligible (long exposure).

**Supplemental Figure 2.**
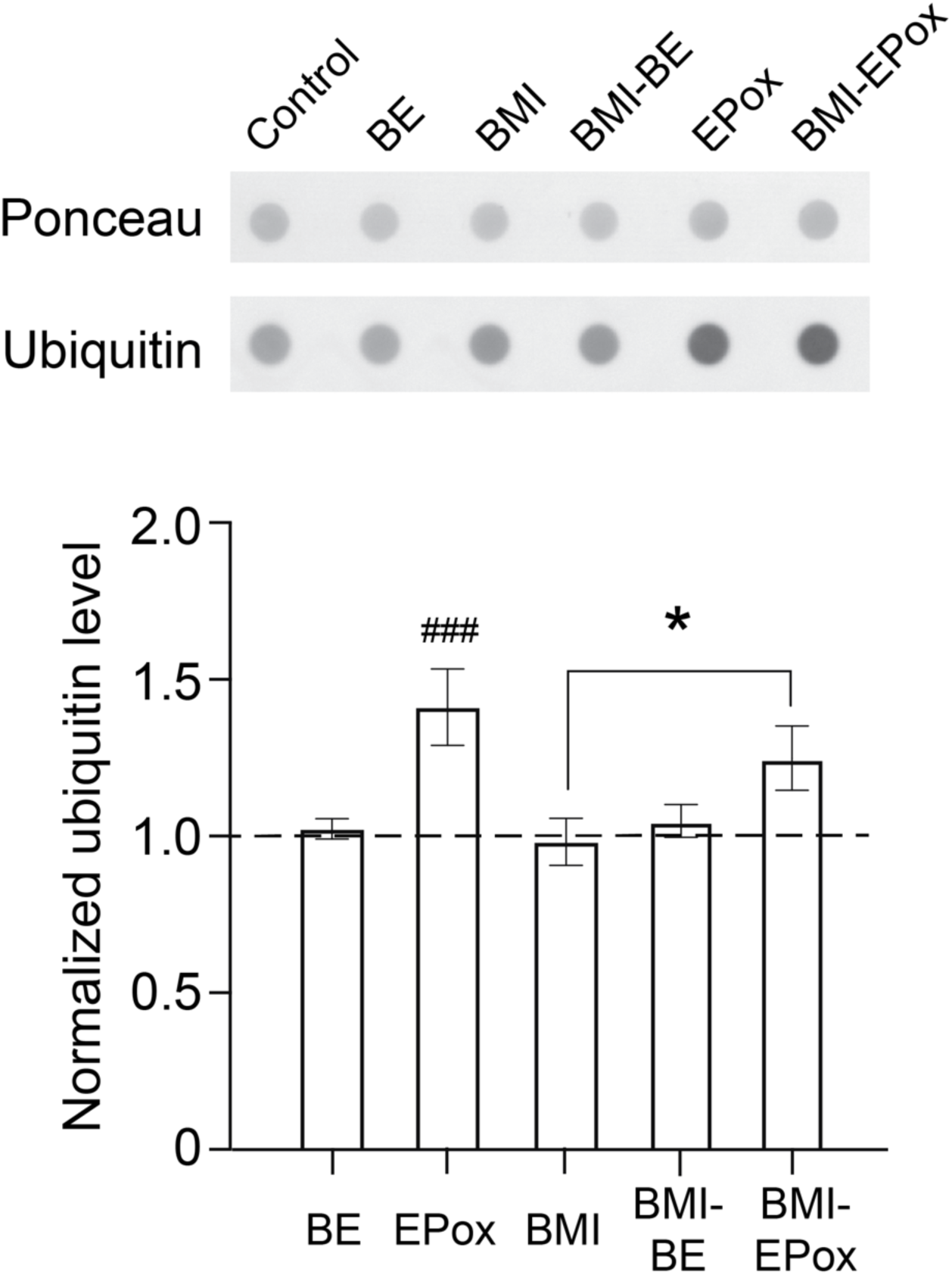
Inhibition of NMPs does not affect ubiquitylation level in the brain tissue. Top: Representative dot blot of total protein samples from animals under basal (control, BE, EPox) or pharmacologically stimulated (BMI, BMI-BE, BMI-EPox) conditions with Ponceau staining and ubiquitin immuno-blotting. Bottom: summary data of quantified ubiquitin level (mean ± s.e.m.). Data from different experimental groups was normalized to the corresponding control group (marked by the dashed line) from the same batch of animals that was run on the same blot. RM one-way ANOVA with Bonferroni’s multiple comparison test. ###: p <0.001, compared to control; *: p < 0.05, comparison as marked, n = 13 experimental batches.

**Supplemental Figure 3.**
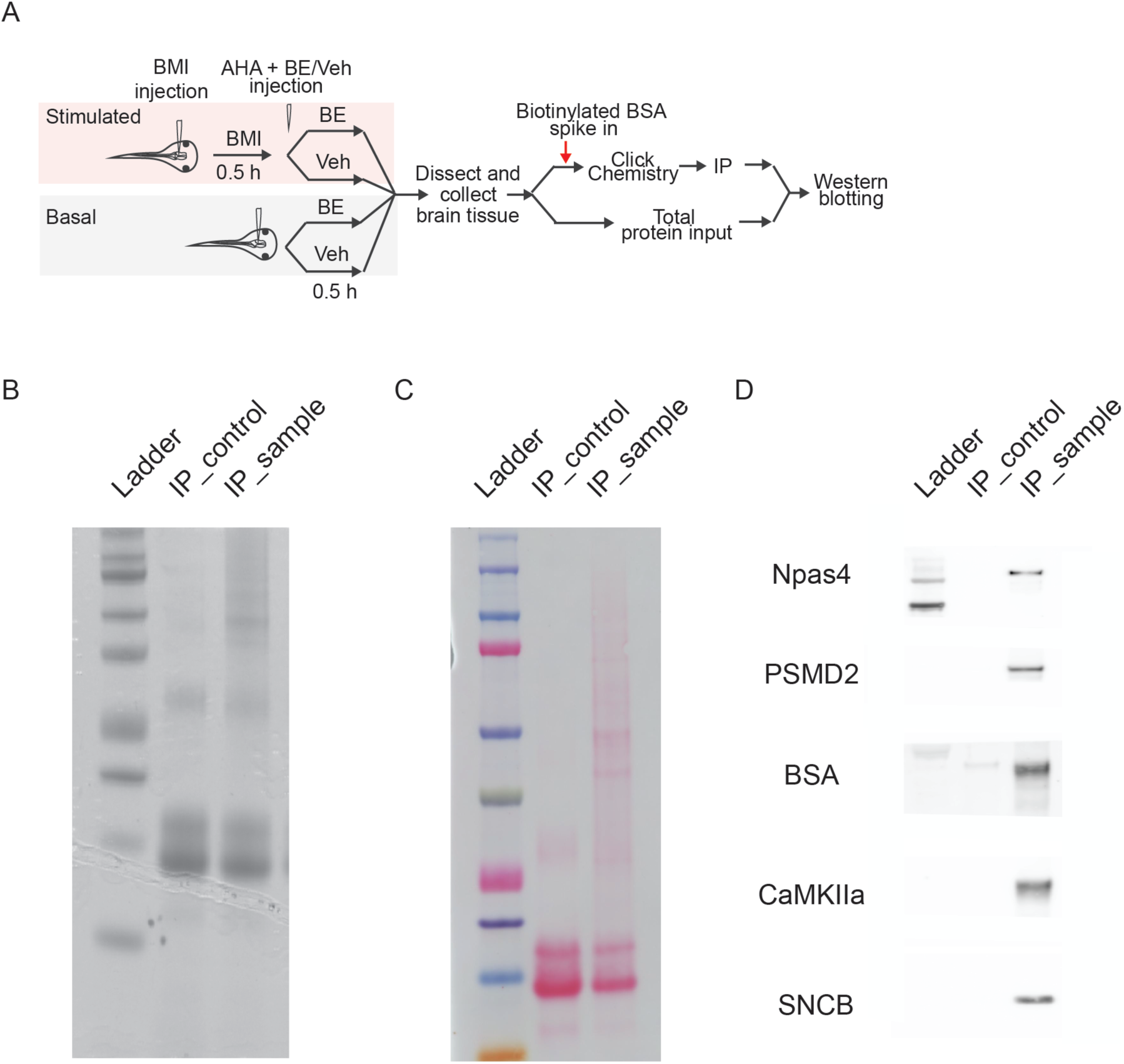
Neutravidin-pull down specifically enriches for AHA-tagged biotinylated nascent proteins. A. Flow chart of experimental protocol (see methods for details). B. Coomassie stained SDS-PAGE gel with eluted samples from the neutravidin-coated agarose resin. IP sample: samples processed with Click chemistry to tag AHA-labelled nascent proteins with biotin molecules. IP-control: same amount of total lysate without Click chemistry to serve as control for the neutravidin pull-down. Very little of protein was seen the IP-control lane suggesting minimum of non-specific binding in the pulldown protocol. The protein bands seen around 15kd and 30kd in the IP-control (and IP) sample was monomers and dimmers of the avidin coating co-eluted from the beads. All samples were prepared from same amount of total lysate input collected from AHA-injected animals. C. Ponceau staining of nitrocellulose blot with eluted samples. D. Western blotting of different individual proteins showed no signal detected in the IP-control sample.

**Supplemental Figure 4.**
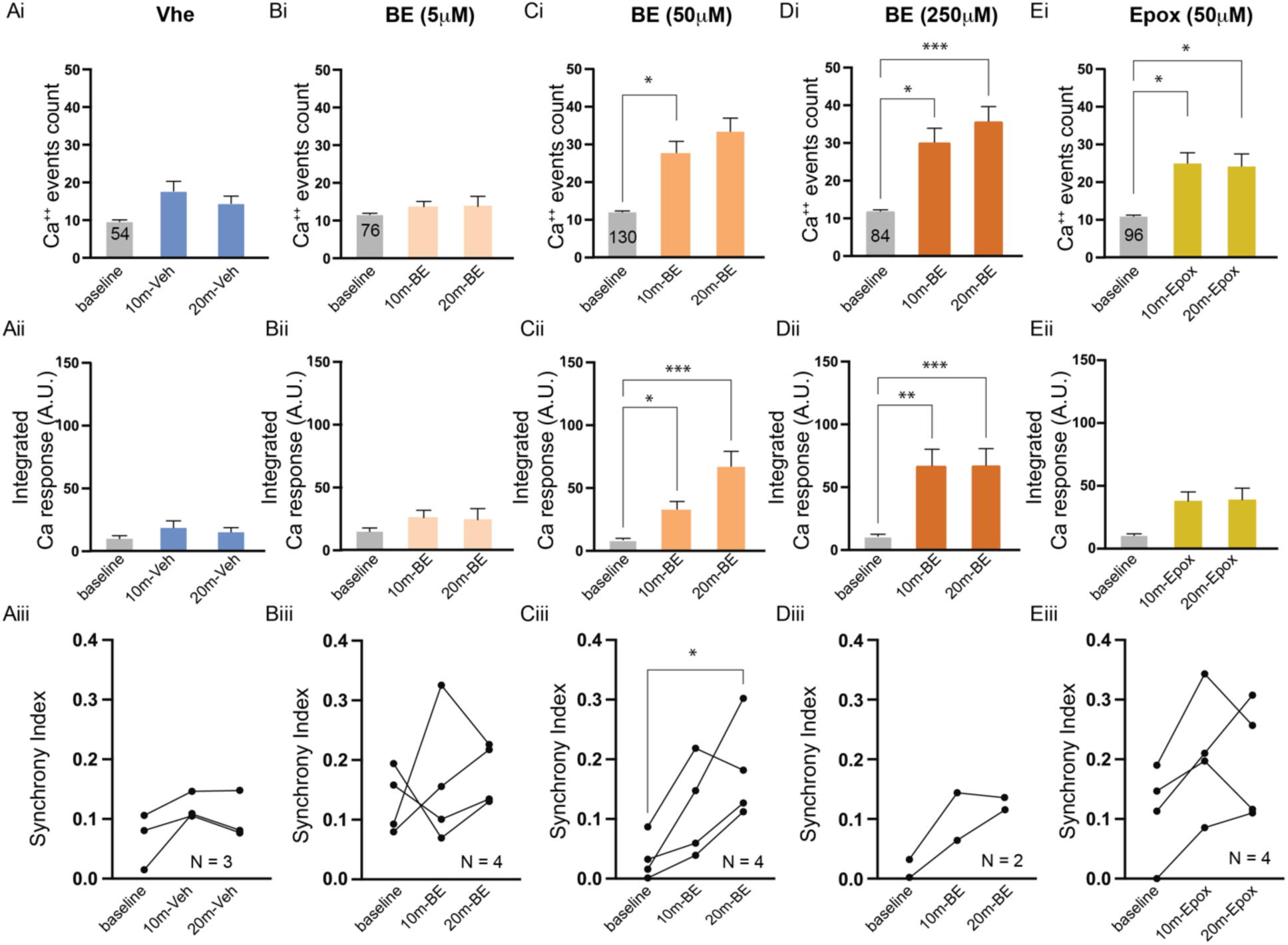
Inhibiting NMP activity increases the spontaneous neuronal activity in tectal neurons under basal condition. A-E. Effects of intraventricular injection of vehicle (Veh, A), Biotin-Epoxomicin (BE), at different injection concentrations (BE 5µM, 50µM, 250µM, B-D), and Epoxomicin (EPox, E) on spontaneous neuronal activity measured by Ca^++^ event counts (i), integrated Ca^++^ response (ii), and network synchrony (iii). n: number of neurons (marked on the histogram). N: number of animals (marked in panel iii). Friedman test with Dunn’s multiple comparison posthoc test. ***: p< 0.001; **: p< 0.01; *: p<0.05.

